# Mapping protein function with CRISPR/Cas9-mediated mutagenesis

**DOI:** 10.1101/076919

**Authors:** Katherine F Donovan, Mudra Hegde, Meagan Sullender, Emma W Vaimberg, Cory M Johannessen, David E Root, John G Doench

## Abstract

CRISPR/Cas9 screening has proven to be a versatile tool for genomics research. We describe a CRISPR/Cas9-mediated approach to mutagenesis, exploiting the allelic diversity generated by error-prone non-homologous end-joining (NHEJ) to identify gain-of-function alleles of the MAPK signaling pathway genes MEK1 and BRAF. These results illustrate a scalable technique to easily generate cell populations containing thousands of endogenous allelic variants of any gene or genes to map variant functions.

---

Deciphering the functional consequences of DNA variation is a defining challenge of the genomic era, and CRISPR/Cas9 technology is the most promising and broadly-developed tool for facile genome engineering^1,2^. Previously, we conducted pooled, genome-wide loss-of-function screens in A375 cells, a melanoma line with the BRAF V600E mutation that is sensitive to MAPK pathway inhibition^3^. These positive selection screens utilized vemurafenib, a BRAF inhibitor, and selumetinib, a MEK inhibitor, to identify sgRNAs that induce drug-resistance in cells and therefore enrich in the cell population over time. As is standard for genetic screens, we then combined information from multiple sgRNAs intended to target the same gene to create a gene-level score. This method identified both previously-validated and novel mediators of this drug resistance phenotype^3,4^. Examination of the sgRNA-level data revealed a curious result, however, namely that in one genome-wide subpool, an sgRNA targeting the gene MAP2K1 (which encodes the protein MEK1) at the site encoding K59 generated the strongest drug-resistance phenotype for both vemurafenib and selumetinib (Table 1). Another MAP2K1 sgRNA screened in a different subpool scored strongly with selumetinib but not vemurafenib. One would expect that sgRNAs targeting MAP2K1, a positive regulator in the pathway inhibited by these drugs, for gene knockout would impair the viability of cells, not rescue them from the drug. Indeed, the other sgRNAs targeting this gene did not lead to drug resistance and were instead strongly depleted (Table 1). We hypothesized that this unexpected result was the consequence of NHEJ-mediated repair of the sgRNA cut site, which led to the creation of drug resistant variants of MEK1.

**Table 1.**
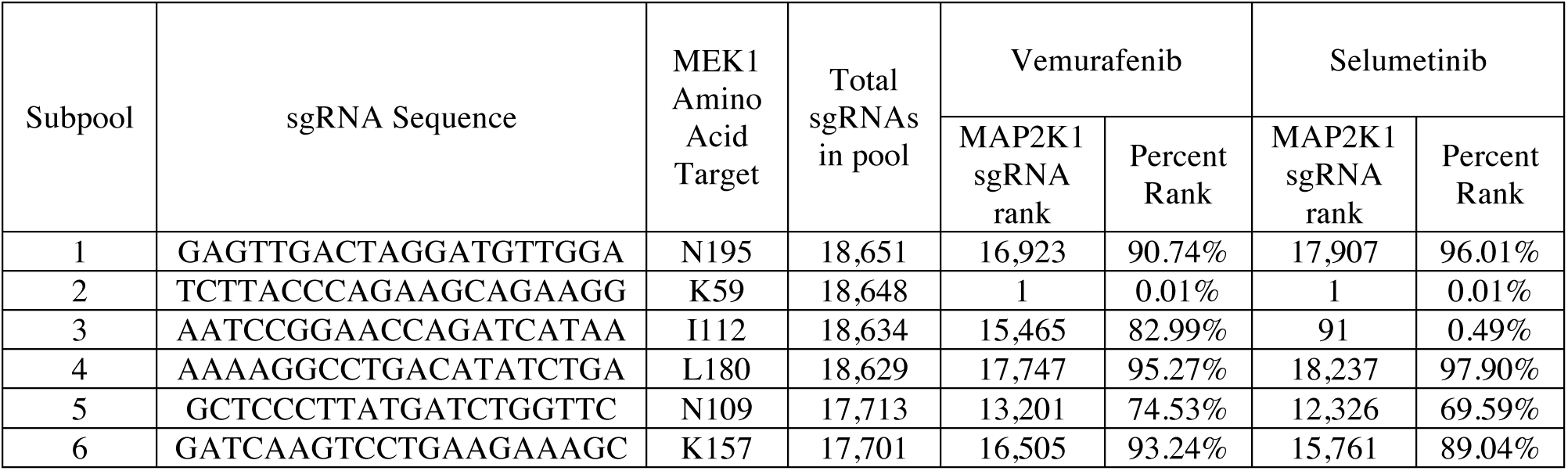
Performance of sgRNAs targeting MAP2K1 in drug resistance screens in A375 cells, from the average of 3 biological replicates^3^.

To test this hypothesis, we introduced the sgRNA targeting MAP2K1 at K59 or a control sgRNA targeting EGFP into A375 cells, treated with either vemurafenib or no selection for two weeks, harvested genomic DNA, and performed PCR with primers flanking the MAP2K1 target site followed by next-generation sequencing. In cells treated with the control sgRNA targeting EGFP, 85% of the MAP2K1 reads mapped perfectly to the wild-type sequence; the majority of the non-matching reads likely represent sequencing errors, as alignment to the reference sequence shows that a majority of the 249 variants present at 25 reads per million (RPM) or greater are single base mismatches (Supplementary Fig. 1). In contrast, for cells treated with the MAP2K1 sgRNA in the absence of selection, only 38% of the reads aligned to wild-type, and there were 1,669 variants present at 25 RPM or greater, coding for 1,351 unique MAP2K1 amino acid sequences. Importantly, many of these variants represent insertions and deletions (indels) of numerous sizes located at multiple positions, results that are consistent with other reports that have examined sgRNA repair products via next-generation sequencing (Supplementary Fig. 2)^4–6^. These results illustrate the large amount of diversity that can be generated by an individual sgRNA.

We next compared allele abundance between vemurafenib-treated and unselected cells, and observed greater than 100-fold enrichment for multiple variants (Fig. 1a). While it is difficult to distinguish true variants from sequencing errors to define an absolute number of variants in a population, we observed that some variants with 18 RPM or fewer in the unselected population were still able to enrich to 1,000 RPM or greater in the vemurafenib-treated population, indicating that they are functionally distinct variants, not sequencing artifacts, and supporting the notion that an individual sgRNA can generate hundreds of alleles.

**Figure 1.**
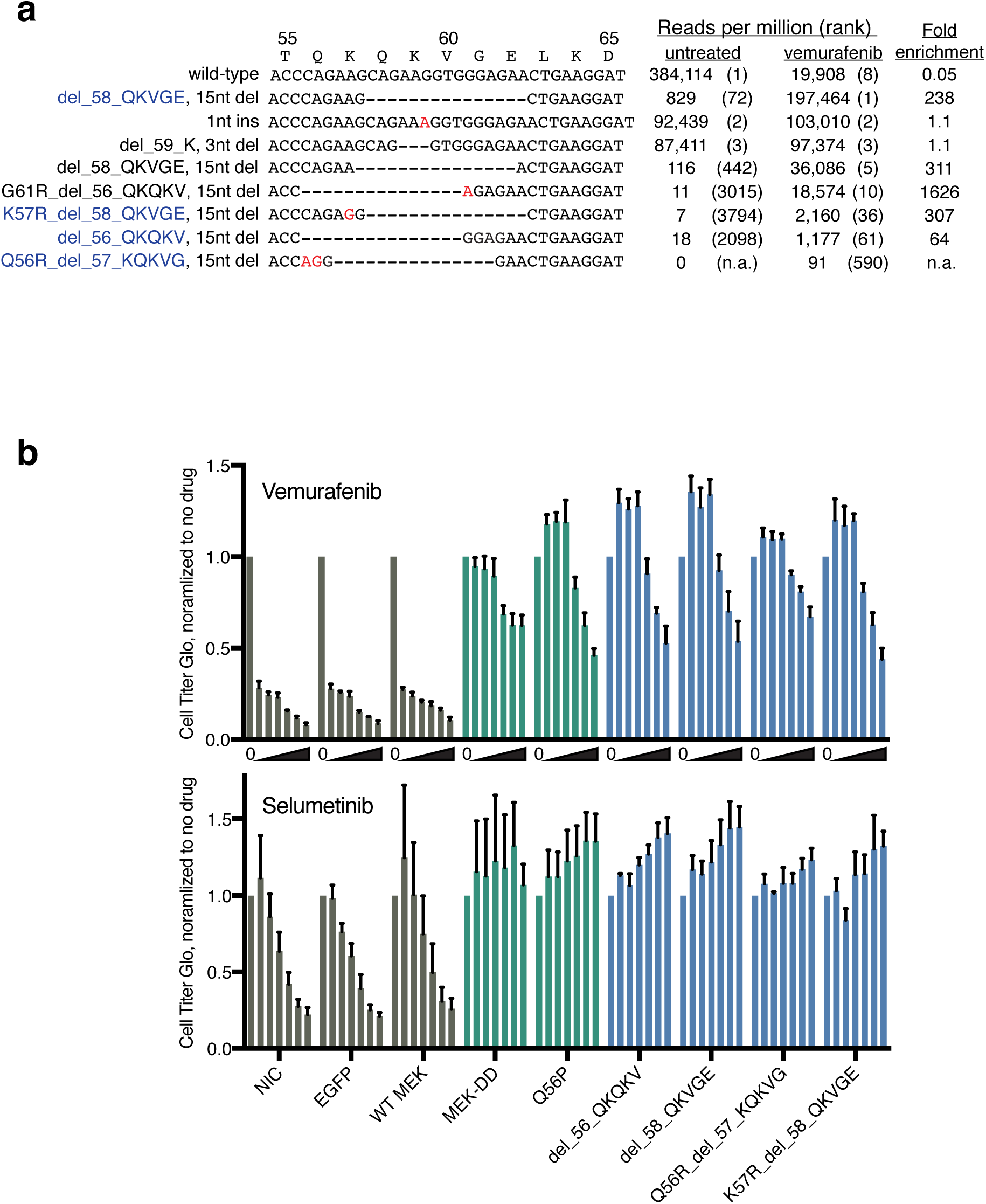
Identification of gain-of-function MAP2K1 alleles. (**a**) Alignment and abundances of MAP2K1 alleles generated in untreated or vemurafenib-selected A375 cells by an individual sgRNA targeting MAP2K1 at the site encoding K59. Mismatched and inserted nucleotides are shown in red. Variants that are studied in the next panel are labeled in blue. (**b**) Cell viability assay for resistance to vemurafenib and selumetinib upon overexpression of ORFs identified in (a). A375 cells constitutively expressing lentiviral-delivered ORFs were assessed for ATP levels after 96 hours of drug treatment, normalized to no drug. Negative controls are shown in gray, positive controls in green, and test ORFs in blue. For vemurafenib, the doses used were, left to right, 0, 0.5, 1, 2, 6, 8, and 10 µM. For selumetinib, the doses used were, left to right, 0, 0.05, 0.1, 0.200, 0.600, 1, and 2 µM.

Many of the variants strongly enriched in vemurafenib contained five amino acid deletions. The amino acids in this region form an alpha-helix that functions to limit the phosphotransferase activity of MEK1^7^; indeed, a study that used *E. coli-*mediated open-reading frame (ORF) mutagenesis followed by overexpression in A375 cells found that a Q56P mutation that disrupts an alpha-helix led to a constitutively-active MEK1^8^. To validate the activity of the variants produced by the K59 sgRNA, we synthesized ORFs with these deletions, introduced them into A375 cells via lentivirus, and treated with selumetinib and vemurafenib. The magnitude of the resistance conferred was as strong as both the well-characterized constitutively-active MEK-DD variant^9^ and the Q56P variant, whereas overexpression of wild-type MEK1 did not produce appreciable resistance (Fig. 1b, Supplementary Fig. 3). These results show that CRISPR/Cas9-mediated mutagenesis of endogenous alleles coupled with strong positive selection can be used to discover novel gain-of-function protein variants and provide insight into regulatory domains.

We next expanded this technique to generate variants across the length of a gene by creating a tiling pool of all possible sgRNAs targeting MAP2K1 (n = 217) and BRAF (n = 279), as well as 100 non-targeting control sgRNAs. We screened this library in triplicate in A375 cells for sgRNAs that confer resistance to vemurafenib or selumetinib compared to untreated cells.

Several sgRNAs targeting MAP2K1 enriched significantly with both drugs, including the original K59-targeting sgRNA, a second sgRNA that also targets K59, and several others that target nearby (Fig. 2a). Interestingly, some sgRNAs generated selumetinib resistance but not vemurafenib resistance. We hypothesized that sgRNAs that led to only selumetinib resistance were disrupting selumetinib binding to MEK1, rather than producing constitutively active mutants. Consistent with this notion, two selumetinib-specific sgRNAs target sites encode V211 and S212, both of which are located in the binding pocket for this class of allosteric inhibitors^7,8^. Additionally, another sgRNA target site encodes I111 in helix C, immediately adjacent to the binding pocket^7,8^. Consistent with this, we previously identified a different sgRNA targeting I112 in a genome-wide screen that induced resistance to selumetinib but not vemurafenib (Table 1), although this individual sgRNA did not score in the tiled library. These results suggest that this method could offer a generalizable approach, complementary to existing techniques, to understanding protein:small-molecule interactions with endogenous proteins expressed from their native promoters. In future work, it will be interesting to compare the spectrum of sgRNAs enriched by selection with a panel of distinct small molecules targeting the same protein that either vary slightly in their chemical composition or target different areas of the protein to determine if this technique is sensitive to such differences.

**Figure 2.**
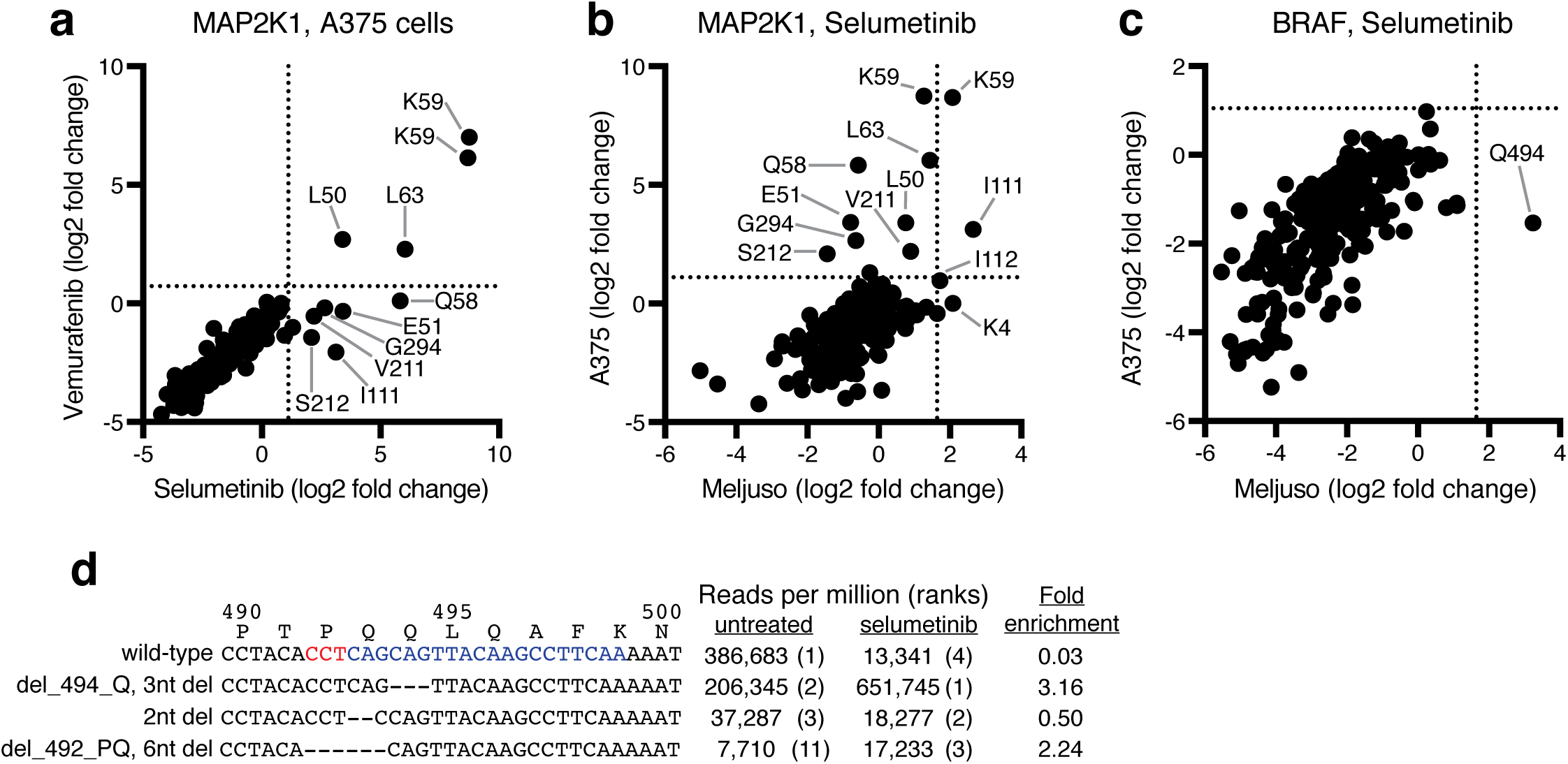
Screening of a pooled sgRNA library tiling the entire length of MAP2K1 and BRAF. (**a**) Comparison of MAP2K1 variants enriched by vemurafenib and selumetinib in A375 cells. The average enrichment across three replicates is shown and dotted lines indicate two standard deviations from the mean of 100 non-targeting sgRNA. (**b**) Comparison of selumetinib-resistant variants of MAP2K1 between A375 and MEL-JUSO cells. The average enrichment across three replicates is shown and dotted lines indicate two standard deviations from the mean of 100 non-targeting sgRNA. (**c**) Comparison of selumetinib-resistant variants of BRAF between A375 and MEL-JUSO cells. The average enrichment across three replicates is shown and dotted lines indicate two standard deviations from the mean of 100 non-targeting sgRNA. (**d**) Alignment and abundances of BRAF alleles generated in untreated or selumetinib-selected MEL-JUSO cells by an individual sgRNA targeting BRAF at the site encoding Q494. The sequence of the sgRNA and PAM are shown in blue and red, respectively.

This same tiling library was also screened for selumetinib resistance in MEL-JUSO cells, a melanoma line carrying the NRAS Q61L oncogene. For MAP2K1, I111- and K59-targeting sgRNAs were enriched, but many other sgRNAs that scored in A375 cells failed to provide significant enrichment in MEL-JUSO cells (Fig. 2b). Conversely, while no BRAF-targeting sgRNAs enriched significantly in A375 cells, one sgRNA targeting Q494 enriched in MEL-JUSO cells (Fig. 2c). Using the same sequencing procedure as described above for the individual MAP2K1-K59 sgRNA, we analyzed the spectrum of alleles generated by this BRAF-Q494 sgRNA in MEL-JUSO cells. In the absence of selection, 1,353 unique variants were present at 25 reads per million or greater. The most-abundant allele in selumetinib-treated cells, accounting for 65% of all sequencing reads, contained a 1 amino acid deletion of Q494, which resides in alpha-helix C, although the overall enrichment for this allele was modest (3.2 fold) as it was already highly abundant in the unselected population (Fig. 2d). The region immediately upstream has been implicated in drug resistance clinically, and structural studies have shown that a 5 amino acid deletion of residues 486 to 490 (∆NVTAP) locks the kinase in an active form^10^. Further work will be necessary to determine if the Q494 deletion has a similar effect on protein structure. These results emphasize the impact of cellular context on variant characterization; both A375 and MEL-JUSO cells harbor activating mutations in the RAS/RAF/MEK signaling cascade. However, these distinct contexts facilitated the identification of MAP2K1 variants in A375 cells and BRAF variants in MEL-JUSO cells.

The results presented here illustrate an experimental strategy that relies on pooled screening, error-prone NHEJ, and strong positive selection to exploit the diversity of variants resulting from CRISPR/Cas9-mediated mutagenesis to map functional domains of proteins. This approach complements existing techniques and offers several advantages. The creation of libraries of variant ORFs either by error-prone PCR, by mutagenesis in *E. coli,* or by synthesis and assembly all rely on exogenous expression of the resulting pool of variants in the cell of interest, whereas the technique described here mutagenizes the endogenous alleles^1,11,12^. Additionally, the former techniques generally create single amino acid substitutions, which may better model mutations that arise from errors in mismatch-repair, while this approach uses each sgRNA to create a family of indels of different sizes and positions, reflective of the heterogeneous outcomes of double-strand DNA break repair. Thus they sample from different areas of sequence space and may reflect different underlying diversity in cell populations.

The approach described here uses relatively small libraries of several hundred unique perturbations for an average-sized gene, relying on the endogenous cellular machinery to generate diversity, while saturating substitution libraries create the diversity *in vitro*, resulting in libraries of thousands of unique constructs. For example, for an average-sized gene encoding a 350 amino acid peptide, all single amino acid substitutions correspond to a library of 6,650 variants. In contrast, the sgRNA library approach described here targets about once every 8 nucleotides – a total of 131 perturbations for a 350 amino acid gene, a 50-fold reduction in library size – and then relies on the cell to generate the variants by NHEJ. As we show, an individual sgRNA can generate ~1,000 variants, meaning that we are testing a library of cells with ~131,000 potential variants, a 20-fold increase relative to a single amino acid substitution library. One useful experimental approach might therefore be to first screen a protein of interest with a tiling sgRNA library as described here, and use the results of that screen to nominate regions for further study by these complementary techniques.

Recently, a CRISPR/Cas9 approach that relies on homology-directed repair (HDR) for mutagenesis has been described, and has the advantage of being able to program the variants of interest^13^. But HDR is a low efficiency process in most cell types, and this technique is limited to cells that can efficiently uptake donor DNA templates, whereas the technique described here is applicable to any cell that can be infected with lentivirus. Another possibility is the use of nuclease-dead Cas9 with DNA modifying domains appended, such as the recently described Base Editor series, although at present only a fraction of potential base changes are accessible by this approach^14^. All Cas9-mediated mutagenesis approaches used to-date are limited in their targeting space by the protospacer adjacent motif (PAM) of NGG required by the *S. pyogenes* Cas9, but the development of alternative Cas9 and related proteins promises to significantly relieve this constraint^15^.

This study began with a serendipitous observation from a genome-wide pooled screen. While the majority of high-scoring sgRNAs in positive selection screens that are not corroborated by other sgRNAs targeting the same gene are likely to be jackpot events or off-target effects, these results highlight the potential to create gain-of-function rather than loss-of-function alleles that, if diagnosed, can be a feature and not a bug.

## METHODS

### Cell line maintenance

Prior to screening, cell lines were maintained without added antibiotics; penicillin/streptomycin was added at 1% during screening. A375 and MEL-JUSO cells were obtained from the Cancer Cell Line Encyclopedia and routinely tested for mycoplasma contamination. Both cell lines were cultured in RPMI 1640 (Invitrogen) supplemented with 10% FBS (Sigma-Aldrich) in a 37°C humidity-controlled incubator with 5.0% CO_2_. Cells were maintained in exponential phase growth by passaging every 2 or 3 days.

### Tiled library design

The MAP2K1/BRAF pooled mini-library included every sgRNA with an NGG PAM along the entire coding sequence and extending slightly into introns of these two genes. Any sgRNAs containing BsmBI restriction sites were excluded to avoid aberrant cleavage during library production. We also added 100 non-targeting control sgRNAs.

### Library production

Pooled library production was performed as described, with oligonucleotides purchased from CustomArray. Briefly, the sgRNA library was digested with Esp3I (Fisher Scientific) and inserted into the lentiCRISPRv2 (Addgene 52961). ElectroMAX Stbl4 electrocompetent cells (Fisher Scientific) were transformed and grown on agar plates. After overnight growth, colonies were harvested and plasmid DNA was isolated and prepped using the Qiagen HiSpeed Maxi kit according to manufacturer’s instructions. The library composition was evaluated by Illumina sequencing.

### Lentivirus production

HEK293T cells were seeded in 9.6 cm^2^ 6-well dishes at a density of 1.5 x 10^6^ cells per well (2mL culture volume) 24 hours before transfection. Transfection was performed using the transfection reagent TransIT-LT1 (Mirus) according to the manufacturer’s protocol. In brief, one solution of Opti-MEM (Corning, 66.25μL) and LT1 (8.75μL) was combined with a DNA mixture of the packaging plasmid pCMV_VSVG (Addgene 8454, 1250 ng), psPAX2 (Addgene 12260, 1250 ng), and the sgRNA library in the transfer vector (plentiCRISPRv2, 1250 ng). The two solutions were incubated at room temperature for 20-30 minutes, during which time the media was changed on the HEK293T cells. After this incubation, the transfection mixture was added dropwise to the surface of the HEK293T cells, and the plates were centrifuged at 1000x*g* for 30 minutes. Following centrifugation, plates were transferred to a 37°C incubator for 6-8 hours, then the media was removed and replaced with media supplemented with 1% BSA. Virus was harvested 36 hours after this media change.

### Pooled screening

#### Determination of viral transduction conditions

In order to determine the amount of virus to use for transduction of the sgRNA library, the virus was titered using the same conditions to be used in the screen. Each cell line was infected in 12-well plates with 100, 200, 300, 500, and 800 μL virus with 3.0 x 10^6^ cells per well. Polybrene was added at 1μg/μL for A375 and 4 μg/μL for MEL-JUSO. The plates were centrifuged at 640x*g* for 2 hours then transferred to a 37°C incubator for 4-6 hours. After this time, each well was trypsinized and split equally by volume to two wells of a 6-well plate. Two days after infection, puromycin was added to one well for each pair seeded post-transduction to select for infected cells expressing the puromycin resistance gene contained in lentiCRISPRv2. After 5 days, cells were counted to determine the efficiency of transduction, represented by the survival of selected cells relative to unselected cells. The virus volume that gave 30 – 50% infection efficiency, corresponding to an MOI of ~0.5 – 1.0, was used for subsequent screening.

#### Screening

The sgRNA MAP2K1/BRAF mini-library was screened in three biological replicates in each of two cell lines. The cells were transduced as described, and two days post-transduction, cells were selected with puromycin (1μg/mL for A375 and 2 μg/mL for MEL-JUSO). Throughout the screen, cells were split at a density to maintain a representation of ~500 cells per sgRNA. Due to the strong positive selection that occurs in the presence of the small molecules, however, some populations fell below this bottleneck; in this case all surviving cells were reseeded at each passage.

After 5 days of puromycin selection, a sufficient number of cells to preserve the initial library representation were seeded for each treatment. Vemurafenib (PLX-4032, Selleckchem S1267) was screened at 2μM, and selumetinib (AZD-6244, Selleckchem S1008) was screened at 1.5μM. An arm with no drug selection was maintained in parallel. Cell counts were taken at each passage to monitor growth. After 14 days in the presence of these small molecules, cells were pelleted by centrifugation, resuspended in PBS, and frozen promptly.

#### Genomic DNA preparation

Genomic DNA (gDNA) was isolated using the QIAamp Blood Mini Kit (Qiagen 51106) as per the manufacturer’s instructions. Briefly, thawed cell pellets were lysed, and genomic DNA was bound to a silica membrane. After two washes, the DNA was eluted using the kit’s Buffer AE. The concentration of these preparations was determined by UV spectroscopy (Nanodrop).

#### PCR Amplification and Next-Generation Sequencing

PCR was used to amplify the sgRNA and to append Illumina flow cell adaptors along with a unique short DNA barcode to allow the samples to be pooled for sequencing while retaining a means to determine which sequence reads arose from which sample, as described previously. All samples were purified by SPRI purification (Agencourt AMPure XP, Fisher Scientific A63880) and then sequenced on a HiSeq 2500 High Output sequencer (Illumina).

#### Tiling pool screen analysis

In the raw results of next-generation sequencing, reads were counted by searching for the CACCG prefix that appears at the 5’ end of all sgRNA constructs in the library. The 20-nucleotide sgRNA insert following this search prefix was mapped to a reference file containing all possible sgRNAs in the library. The read was then assigned to a sample by cross-referencing with a conditions file indicating which sample (i.e. a single well on the PCR plate) corresponded to each barcode. The resulting matrix was then normalized to reads per million (RPM) by the following calculation: reads per sgRNA/total reads per condition x 10^6^. This value was then log_2_-transformed after first adding 1 read to each sgRNA to eliminate zero values, and this final value was denoted as the lognorm. Enrichment of sgRNAs was determined using the log_2_ fold-change for each sgRNA relative to its abundance in the original plasmid DNA (pDNA) of the library. This is calculated by first averaging replicates within the PCR plate, then subtracting the lognorm for each sgRNA in the pDNA from the same sgRNA in the screen endpoint gDNA.

### Analysis of variants produced by individual sgRNAs

#### Infections

For sgRNAs that showed strong enrichment under small molecule selection, single-sgRNA infections were performed to validate the screening result and to reveal via deep sequencing the specific indels formed by CRISPR/Cas9-mediated cutting. Virus was prepared for each sgRNA as described, and transductions were performed in A375 and MEL-JUSO cells as described using a virus volume determined to transduce at least 1 x 10^6^ cells (300 μL for both A375 and MEL-JUSO). Cells were selected for 5 days with puromycin (1μg/mL for A375 and 2 μg/mL for MEL-JUSO) to remove uninfected cells, then each population of transduced cells were split to vemurafenib (2 μM), selumetinib (1.5 μM), and unselected arms. To maintain representation of the library, at least 1 x 10^6^ cells were reseeded with each passage. Due to the rare occurrence of gain-of-function variants, however, some populations fell below this bottleneck; in this case all surviving cells were reseeded at each passage. After 14 days in the presence of these small molecules, cells were pelleted by centrifugation at 1000x*g* for 5 minutes, resuspended in PBS, and frozen promptly.

#### PCR Amplification

For MAP2K1, PCR primers included all sequences necessary for Illumina sequencing were designed to amplify across the sgRNA cut-site. For BRAF, two rounds of PCR were performed: a first round amplified across the cut site, and second round appended the necessary Illumina sequences. PCR-amplified samples were purified using SPRI, and samples were sequenced on a MiSeq desktop sequencer (Illumina).

### MAP2K1

Fwd: AATGATACGGCGACCACCGAGATCTACACTCTTTCCCTACACGACGCTCTTCCGATCTCTAGAGCTTGATGAGCAGCA

Rev: CAAGCAGAAGACGGCATACGAGATGACCTTAGGTGACTGGAGTTCAGACGTGTGCTCTTCCGATCTCTCACTGATCTTCTCAAAGT

### BRAF

Fwd: TTGTGGAAAGGACGAAACACCG GAGACTTGGAGTAACAATTGCC

Rev: TCTACTATTCTTTCCCCTGCACTGT CCACTGGGAACCAGGAGC

#### Sequencing Analysis

All sequencing reads were first assessed for on-target amplification by requiring an exact match of the 10 nucleotide string that appears immediately downstream of the primer binding site in the genomic DNA target site. The next 62 (MAP2K1) or 111 (BRAF) nucleotides were then analyzed and the number of reads for each unique sequence was counted and then normalized to reads per million (RPM).

#### Variant Analysis

BLAST was used to align sequences to the MAP2K1 reference sequence for the samples treated with sgRNA-MAP2K1 (n = 1,669) and sgRNA-EGFP (n = 249) variant sequences with at least 25 RPM (**Supplementary Table 2**). The parameters used for the alignment were a word size of 11, gap open cost of 5, gap extend cost of 2, nucleotide match reward of 2, and nucleotide mismatch penalty of -3. The btop (Blast traceback operations) string was used to parse the count and positions of mismatches, insertions, and deletions for each variant. Sequences were further analyzed (Supplementary Figs 1, 2) if the alignment extended for at least 59 nucleotides, including gaps, which resulted in 246 sequences for sgRNA-EGFP and 1,241 sequences for sgRNA-MAP2K1.

### ORF over-expression

Open reading frames (ORFs) of wild-type MEK1 and mutants thereof were synthesized (Genscript) and cloned into the lentiviral vector pLX_304 (Addgene 25890)^16^. For each construct, virus was prepared as described above. A375 cells were seeded at 30,000 cells/mL in a 96 well plate, and infected in triplicate with 5 μL of each ORF virus in the presence of polybrene (5μg/mL). The vector pRosetta, which expresses EGFP (Addgene 59700), was used as a transduction control. The MEK-DD construct was described previously and cloned into pLX_304^17^. After addition of the viruses, the plates were centrifuged for 30 minutes at 230xg. After 2 days of incubation, vemurafenib or selumetinib were added to each plate over a range of doses. After 4 days in the presence of these small molecules, cell viability was determined using the CellTiterGlo Luminescent Cell Viability Assay (Promega G7573) according to manufacturer’s instructions. Luminescence was detected and quantitated on the EnVision plate reader (Perkin Elmer).

## ACKNOWLEDGEMENTS

We thank T. Mason and the Walk Up Sequencing team for Illumina sequencing support; M. Tomko, M. Greene, A. Brown, D. Alan, and T. Green for software engineering support; O. Bare and S. Amaral for operations support; S. Milczarek and X. Yang for library production support (Broad Institute). J.G.D. is a Merkin Institute Fellow and is supported by the Next Generation Fund at the Broad Institute of MIT and Harvard.

---

**Supplementary Figure 1.**
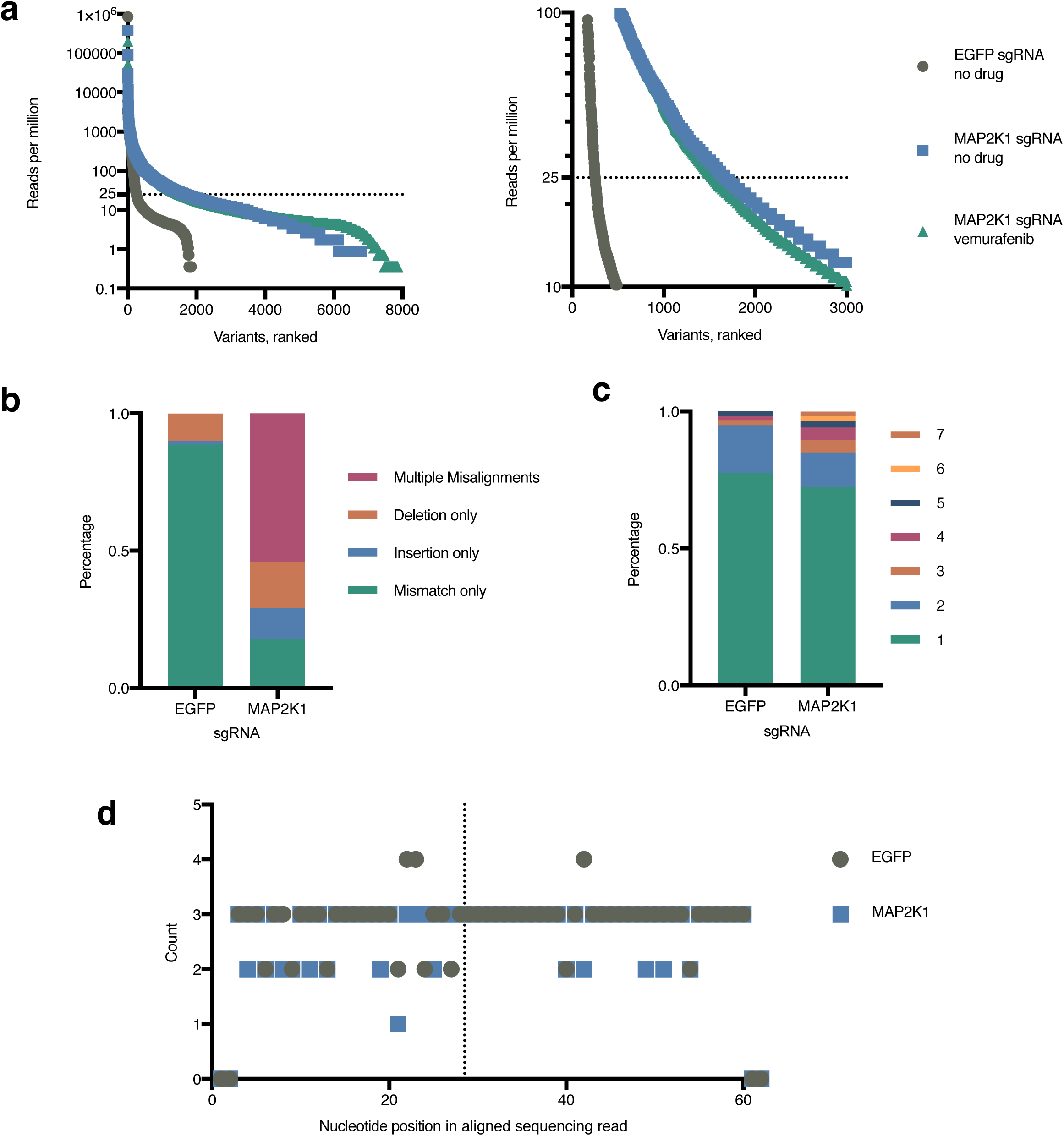
An individual sgRNA generates many allelic variants. (**a**) Within each of three conditions, variants detected by deep sequencing of the MAP2K1 locus were ranked by their abundance and the reads per million is plotted. Left panel shows all the variants, right panel enlarges one region. (**b**) For the two sgRNAs in the absence of drug selection, all reads present at an abundance of 25 reads per million (RPM) or higher were aligned to the wild-type reference. Those which were able to align were classified into one of four categories. (**c**) For the reads in the mismatch-only category from panel b, the number of mismatches is plotted. (**d**) For the reads with only 1 mismatch, the position is plotted. The dotted line indicates the cut site of the sgRNA. Note that BLAST categorizes unaligned nucleotides that occur at either end of the string as deletions

**Supplementary Figure 2.**
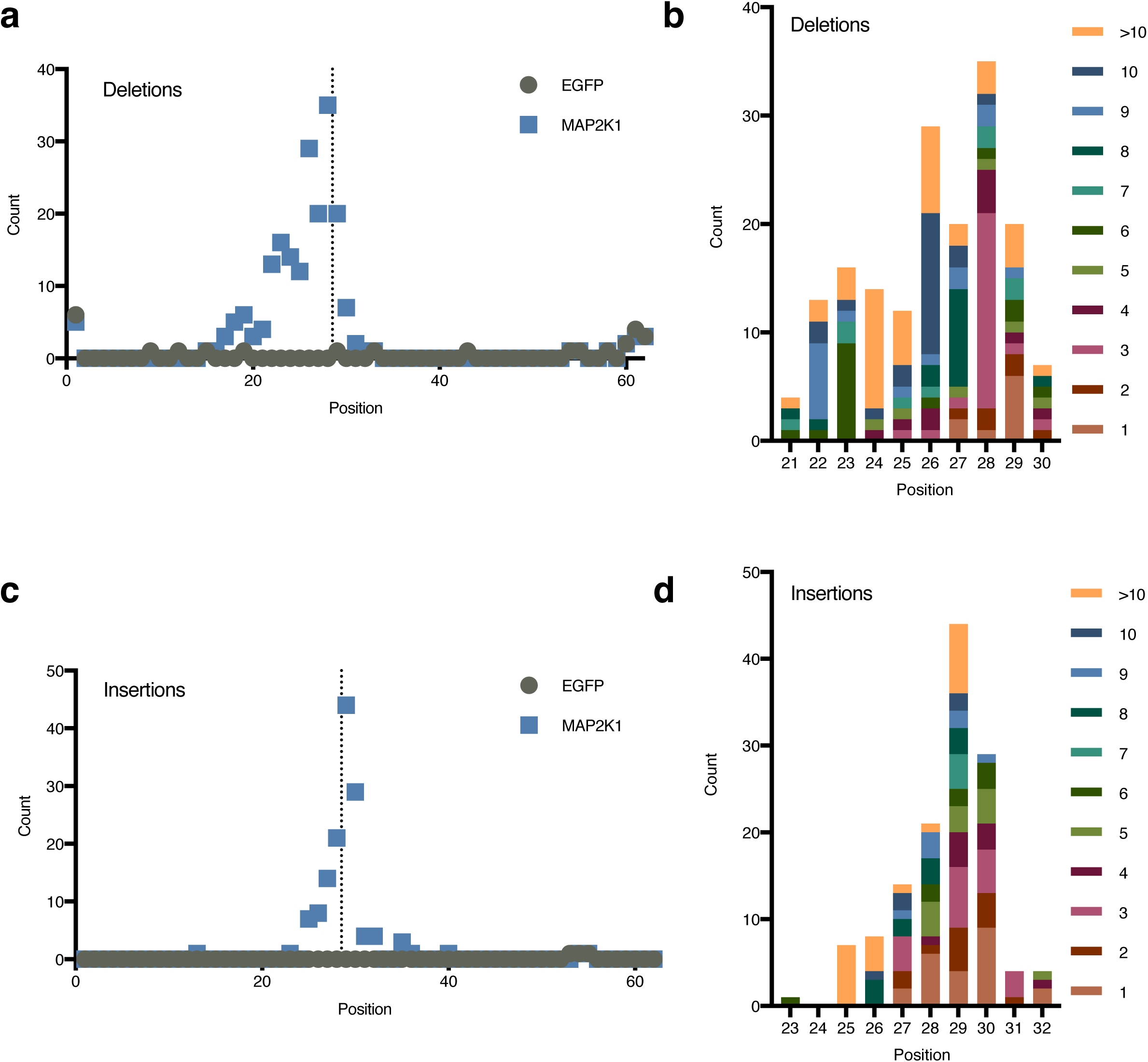
Analysis of indels from deep sequencing of MAP2K1 at the sgRNA target site. (**a**) For all reads present at an abundance of at least 25 reads per million (RPM) and that were categorized as deletion only in Supplementary Figure 1b, the number of unique deletions detected in the MAP2K1 locus for cells treated with the indicated sgRNA and no drug selection is plotted by their position. Note that BLAST categorizes unaligned nucleotides at either end of the string as deletions. The dotted line indicates the position of the MAP2K1 sgRNA cut site. (**b**) For the deletions generated by the MAP2K1 sgRNA, the frequency of deletions categorized by their size is plotted. (**c**) As in panel a, but for insertions. (**d**) As in panel b, but for insertions.

**Supplementary Figure 3.**
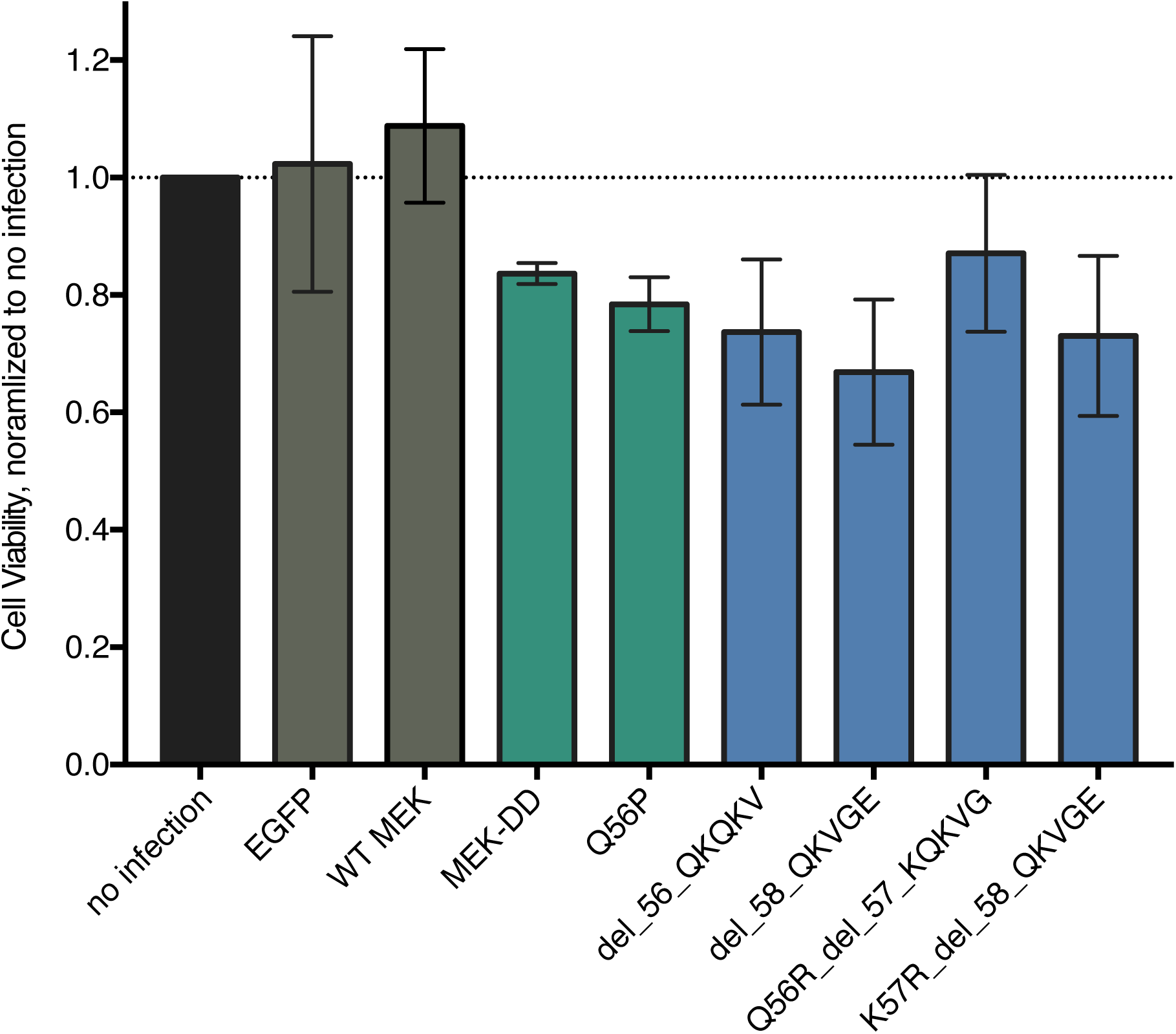
Cell viability upon ORF overexpression of MEK1 variants. A375 cells were infected with lentivirus expressing the indicated ORFs and assessed for cell viability via Cell Titer Glo after 7 days. Error bars represent the standard deviation of triplicate experiments, normalized to the no infection control. Negative controls are shown in gray, positive controls in green, and test ORFs in blue. The MEK1 variants showed a modest growth suppresion in the absence of drug selection.

## REFERENCES

1. Fowler, D. M., Fowler, D. M., Fields, S. & Fields, S. Deep mutational scanning: a new style of protein science. Nature Methods 11, 801–807 (2014).

2. Doudna, J. A. & Charpentier, E. The new frontier of genome engineering with CRISPR-Cas9. Science 346, 1258096–1258096 (2014).

3. Doench, J. G. et al. Optimized sgRNA design to maximize activity and minimize off-target effects of CRISPR-Cas9. Nat Biotechnol 34, 184–191 (2016).

4. Shalem, O. et al. Genome-Scale CRISPR-Cas9 Knockout Screening in Human Cells. Science 343, 84–87 (2014).

5. van Overbeek, M. et al. DNA Repair Profiling Reveals Nonrandom Outcomes at Cas9-Mediated Breaks. Mol. Cell 63, 633–646 (2016).

6. Wang, T., Wei, J. J., Sabatini, D. M. & Lander, E. S. Genetic Screens in Human Cells Using the CRISPR-Cas9 System. Science 343, 80–84 (2014).

7. Fischmann, T. O. et al. Crystal structures of MEK1 binary and ternary complexes with nucleotides and inhibitors. Biochemistry 48, 2661–2674 (2009).

8. Emery, C. M. et al. MEK1 mutations confer resistance to MEK and B-RAF inhibition. Proceedings of the National Academy of Sciences 106, 20411–20416 (2009).

9. Johannessen, C. M. et al. COT drives resistance to RAF inhibition through MAP kinase pathway reactivation. Nature 468, 968–972 (2010).

10. Foster, S. A. et al. Activation Mechanism of Oncogenic Deletion Mutations in BRAF, EGFR, and HER2. Cancer Cell 29, 477–493 (2016).

11. Melnikov, A., Rogov, P., Wang, L., Gnirke, A. & Mikkelsen, T. S. Comprehensive mutational scanning of a kinase in vivo reveals substrate-dependent fitness landscapes. Nucleic Acids Research 42, e112–e112 (2014).

12. Kitzman, J. O., Starita, L. M., Lo, R. S., Fields, S. & Shendure, J. Massively parallel single-amino-acid mutagenesis. Nature Methods 12, 203–206 (2015).

13. Findlay, G. M., Boyle, E. A., Hause, R. J., Klein, J. C. & Shendure, J. Saturation editing of genomic regions by multiplex homology-directed repair. Nature 513, 120–123 (2014).

14. Komor, A. C., Kim, Y. B., Packer, M. S., Zuris, J. A. & Liu, D. R. Programmable editing of a target base in genomic DNA without double-stranded DNA cleavage. Nature 533, 420–424 (2016).

15. Mohanraju, P. et al. Diverse evolutionary roots and mechanistic variations of the CRISPR-Cas systems. Science 353, aad5147 (2016).

16. Yang, X. et al. A public genome-scale lentiviral expression library of human ORFs. Nature Methods 1–6 (2011). doi:10.1038/nmeth.1638

17. Boehm, J. S. et al. Integrative Genomic Approaches Identify IKBKE as a Breast Cancer Oncogene. Cell 129, 1065–1079 (2007).

